# Direct interactions of CEACAM1 and CD36 with micellar LPS and each other

**DOI:** 10.64898/2026.02.01.703119

**Authors:** Weidong Hu, Patty Wong, Supriyo Battacharya, Zhuo Li, Lin Li, Eric Aniogo, Jitender Jitender, Teresa Hong, Zhifang Zhang, Paul Yazaki, Maciej Kujawski, John E. Shively

## Abstract

Micellar lipopolysaccharide (mLPS), a major shed product of gram-negative bacteria in the intestine, may result in TLR4-mediated sepsis in the circulation. CEACAM1 and CD36, expressed on both epithelia and immune cells, regulate TLR4 signaling and inflammation, suggesting a shared role in innate immunity. Furthermore, CEACAM1 associates with CD36 on hepatocytes, regulating lipid storage and bile acids (BAs) in the liver, where LPS detoxification occurs. Using *E. coli* mLPS-Ra as a model and soluble forms sCEACAM1 or sCD36, we assessed their direct binding to mLPS by SPR, SEC, and TEM. SPR derived steady state affinity constants (K_D_= 1.47 x 10^-6^ M and K_D_= 2.23 x 10^-6^ M) were obtained for mLPS-Ra binding to sCEACAM1 and sCD36, respectively. mLPS-Ra binding to sCEACAM1 and sCD36 was reduced by bile acids in order of sodium deoxycholate > sodium cholate. mLPS exhibited dose dependent binding to sCEACAM and sCD36 by SEC and the isolated complexes formed well defined micelles on TEM. A K_D_ of 5.28 x 10^-8^ M was obtained for sCEACAM1 binding to immobilized sCD36. Antibody-based-proximity ligation assays (PLA) demonstrated the association of the ectodomains of CEACAM1 and CD36 on hepatic (P< 0.001) and HEK cells (P< 0.018), while biotin-based PLA demonstrated association of the cytoplasmic domains of CEACAM1 and a CD36-BioID2 fusion protein in HEK cells. Alpha Fold predicted CD36 binding in cis to CEACAM1 in a membrane model, in agreement with the results of the PLAs. Based on the SPR, SEC and TEM binding studies, CEACAM1 and CD36 share a cooperative mLPS binding function distinct from their regulation of TLR4 binding.

## INTRODUCTION

Human CEACAM1 is a single pass transmembrane glycoprotein with 3-4 extra-cellular Ig-like domains and two different cytoplasmic signaling domains formed by alternative mRNA splicing (1). It occurs as inactive cis-dimers on epithelial cells and immune cells, and upon activation by inside-out signaling, forms trans-dimers between cells that modulate a variety of co-receptor signaling pathways, including the TCR of T-cells (1) and the BCR of B-cells (2,3). Dimer formation is dependent on a G-xxx-G motif in its transmembrane domain (4), leading to subsequent homophilic binding of the GFCC’C’’ faces of its distal IgV-like N-domain (5,6). The large size of the *CEACAM* gene family with a conserved N-domain (7) suggests functions for this domain in addition to cell-cell adhesion. Less confirmed studies suggested that the N-domain of CEACAM1 may also interact with bile acids (8), ATP (9), and LPS (10). Importantly, CEACAM1 is highly expressed in bile canaliculi and secreted into bile as a soluble protein known as BGP1 (11-13), and can modulate LPS associated TLR4 signaling (14,15). In addition, *Ceacam1* knock out mice are extremely sensitive to LPS (15) and develop fatty liver and obesity on normal chow (16,17). With these studies in mind, along with high CEACAM1 expression throughout the intestinal tract (18) where LPS is found at high levels, we studied its interactions with LPS and bile acids in more detail.

The heightened interest in the role of the microbiome in gut homeostasis (19), includes studies on the role of intestinal receptors for microbial products such as LPS. Among candidates for this function, the scavenger receptor CD36 stands out for its uptake of LPS in the intestine (20,21), its modulation of TLR4 signaling (22) and its colocalization with CEACAM1 in hepatocyte lipid droplet formation (23). Furthermore, CD36, like CEACAM1 is found as a soluble form in serum and its levels correlate with fatty liver disease (24) in which LPS has a significant role (25,26). Based on these correlations, we included CD36 in our LPS binding studies.

The molecular nature of LPS binding to a number of LPS binding proteins demonstrates the requirement that its hydrophobic acyl chains, numbering from 4-6 in most examples, must be buried in the interior of the protein (27-30). Since micellar LPS (mLPS) is the physiological relevant form of LPS shed from growing and/or dying gram negative bacteria (31,32), the transfer of monomeric units from the ends of mLPS are the most likely mechanism of LPS-protein complex formation. As an example, mLPS, along with fatty acids, is taken up by CD36 in the small intestine (20) where CEACAM1 is also expressed (33,34). Like all intestinally absorbed lipids that enter the lacteal lymphatics, mLPS may be incorporated into circulating lipid particles including chylomicrons, VLDL, LDL and HDL where it does not usually represent a threat to the organism (35,36).

Given these considerations, we hypothesized that CEACAM1 and CD36 both directly bind mLPS, and also, directly interact with each other to regulate mLPS uptake. We tested these hypotheses with direct binding and ultrastructural approaches and molecular models.

## METHODS

### Materials

Antibodies used in these studies are described in the sections below. Lipopolysaccharide (LPS) rough strain from *E. coli* EH 100 (Ra mutant) was purchased from Sigma. Sodium dodecyl sulfate 0.5% (w/v), HBS-N buffer and HBS-EP buffer were purchased from Cytiva. Sodium cholate hydrate and sodium deoxycholate monohydrate were purchased from Thermofisher Scientific. Human recombinant ectodomain CD36 (Gly 30 – Asn 439) was purchased from AcroBiosystems.

### Expression of CEACAM1 proteins

Plasmid construction and membrane expression of CEACAM1-4L has been previously described (37,38). The soluble ectodomain of human sCEACAM1-4D (4 domains) and soluble N-domain of CEACAM1 containing C-terminal His6 tags with its secretion signal sequence were cloned into the mammalian expression vector pIBS54. The constructs were transiently expressed in Expi293 cells using the ExpiFectamine™ 293 Transfection Kit (Thermo Fisher Scientific, Waltham, MA) following the manufacturer’s protocol. Cells were maintained in Expi293 expression medium at 37 °C in a humidified incubator with 8% CO₂ and agitation at 125 rpm. The culture supernatant was collected seven days after transfection.

The harvested medium was clarified by centrifugation and treated overnight at 4 °C with AG1-X8 resin (Bio-Rad, Hercules, CA) to remove residual nucleic acids. The treated supernatants were diluted with wash buffer to get 10mM imidazole concentration then applied to a 5 ml Ni-NTA Superflow column (Qiagen, Germantown, MD) pre-equilibrated with buffer containing 50 mM sodium phosphate, 300 mM sodium chloride, and 10 mM imidazole at pH 7.4. The column was washed with the same buffer containing 20 mM imidazole, and bound protein was eluted with 250 mM imidazole. The eluate was buffer exchanged into 0.05 M MES, 0.01 M potassium phosphate buffer, pH 6.5, using 10 kDa molecular weight cutoff centrifugal filters. The IMAC-purified proteins were further polished using ceramic hydroxyapatite (CHT) Type I (Bio-Rad, 20 µm, 7 mL) chromatography to remove aggregates. The column was equilibrated with 0.05 M MES, 0.01 M potassium phosphate buffer at pH 6.5. After loading, the column was washed with 0.05 M MES, 0.05 M potassium phosphate buffer, and the proteins were eluted using a linear gradient up to 0.2 M potassium phosphate. Protein elution was monitored by absorbance at 280 nm. Monomeric fractions were confirmed by size-exclusion chromatography (Superdex 200 Increase 10/300 GL, Cytiva) and exchanged into PBS for subsequent experiments

### Surface Plasmon Resonance (SPR)

The experiments were carried out on a Biacore^TM^ X100 (GE Healthcare) at 25°C. Human recombinant human CD36 (CD6-H5221 from Acro Biosystems) was diluted into 10 mM sodium acetate (pH 4.5) to final concentration of 12.5 µg/mL. The CD36 was immobilized to two NHS-CM5 chips with resonance units (RU) of ∼430 and ∼2500. The immobilized CD36 with 430 RUs was used to study the interaction with Ceacam1, and the one with 2500 RUs was used to study the interaction with LPS and/or bile acids. The expression of human recombinant sCEACAM1-4D was described above. A concentration of 50 µg/mL of sCEACAM1-4D was prepared in 10 mM sodium acetate (pH 4.5) for immobilization onto a CM5 with RU of ∼2300. Concentration of N-domain of Ceacam1 of 50 ug/ml in 10 mM sodium acetate (pH 4.5) is used for immobilization onto a CM5 chip with RU of ∼1050. The running buffer HBS-EP was used for CEACAM1 interaction with immobilized sCD36, and the chip was regenerated with 0.1 N HCl. The running buffer HBS-N was used for the interaction between LPS or bile acids with immobilized sCEACAM1-4D or sCD36, and the chip was regenerated with 0.25% or 0.5% SDS water solution. Typical contact and dissociation times for binding studies were set at 180 and 700 seconds, respectively. The contact and stabilization times for regeneration were set to 30 and 700 seconds, respectively. The measurements were carried out twice for all concentration points. Before analysis, the data was corrected to reference cell, as well as, to blank buffer readings to reduce the bulk refractive index shift and systematic errors. Data was analyzed and presented using Biacore^TM^ X100 software and Excel. K_D_ values were calculated using both kinetic and steady-state affinity approaches in the Biacore^TM^ X100 software.

### Size exclusion chromatography and calculation of Stoke’s radius

Samples were run on a Superdex 200 Increase 10/300 GL column from Cytiva, Marlborough, MA) in pH 7.4 PBS at a flow rate of 0.5 mL/min with UV absorption at 215 and 280 nm. The column was calibrated with standards of known Stokes radius (BioRad gel calibration standards) and calculations performed according to La Verde et al. (39). The column and HPLC system were rendered endotoxin free by washing with 3 column volumes of 1 N NaOH followed by 6 column volumes of endotoxin free PBS.

### Transmission electron microscopy

A 5 µl drop of protein complexes isolated from SEC was applied to a freshly plasma-cleaned 200-mesh formvar/carbon coated grid (Ted Pella Prod No. 01801) for 1 minute, blotted to a thin film using filter paper, floated on drops of Milli-Q H_2_O for 20 seconds (3x), blotted to a thin film again using filter paper, and immediately stained with 1% (w/v) uranyl acetate for 1 minute. The stain was then blotted away with filter paper and the grid subsequently air-dried. Transmission electron microscopy was performed using an FEI Tecnai 12 electron microscope operating at 120 kV equipped with a Gatan OneView CMOS camera. Images were collected at nominal magnification of 52,000x (0.21 nm/pixel) using Gatan Digital Micrograph. Images were acquired at a nominal under focus of -1.5 μm.

Particle size analysis was carried out using automatic particle counting function in Image J (40). For sCEACAM1-4D, individual particles in the 52,000x images were selected using automated picking protocols, followed by several rounds of reference-free 2D alignment and classification algorithms using XMIPP (41) to sort them into self-similar groups.

### Proximity ligation assays

Proximity ligation assay (Duolink PLA, Millipore Sigma) for CD36 and CEACAM1 was performed according to the manufacturer’s protocol. Briefly, cells were grown in Nunc Lab-Tek II Chamber Slide System (Thermo Fisher), fixed and permeabilized, blocked and incubated with primary antibodies: huCD36 (D8L9T) Rabbit mAb (Cell Signaling Technology, Danver, MA) and huCEACAM1 (MAB22441) Mouse mAb (R&D Systems, Mineapolis, MN) overnight in 4°C. Next day slides were incubated with anti-Mouse and anti-Rabbit PLA probes, ligated and amplified by rolling circle amplification using Phi-X29 polymerase. Slides were mounted with In Situ Mounting Medium with DAPI and analyzed using a LSM900 confocal microscope (Carl Zeiss Microscopy, White Plains, NY) followed by quantitation of signals and statistical analysis using a paired t-test.

Biotin proximity ligation assay. Transient expression of CD36-BioID2 and CEACAM-1 or both was performed using the Expi293F mammalian expression system (Thermofisher, Waltham, MA, USA). Briefly, 75 x 10^6^ viable cells were transfected with 25 μg of CD36-BioID2 and/or CEACAM1-4L plasmids in a 125mL vented, plain bottom flask (PBV125). For co-expression, equal volumes of both CD36-BioID2 and CEACAM-1 plasmids (12.5 μg each) were also transfected, and a flask with no DNA was set as a negative control. On day 4 following transfection, the cells were collected and stained with either anti-CD36 (FAB19551G) or anti-CEACAM-1 (R&D systems, FAB2244P) antibody to assess the level of expression using flow cytometry.

Proximity labelling was performed by adding biotin (Thermo Fisher, B20656) to a final concentration of 50 µM to each group and incubated overnight at 37°C. Next, 20 x 10^6^ cells were transferred to a 15 mL conical tube and were washed twice with 15 ml ambient PBS. Following the wash, the cells were lysed with 2 mL of lysis buffer with 20 μL protease inhibitors. The lysed cells were transferred to a 2 mL microcentrifuge tube and incubated for 30 min on ice. The lysate was centrifuged at 15,000 g for 5 min at 4°C and the clarified supernatant transferred to a new 2 mL tube. The labelled protein was isolated by incubating the lysate with 100 μL of streptavidin beads (Thermo Fisher, 65801D) and incubated on a rotator for 4 hrs at 4°C. Thereafter, the beads were washed three times to remove unbound protein and resuspended with 400μl of SDS-PAGE sample buffer for Western blot analysis.

Western blot analysis was done with a detection protocol using the Quick Western Kit (Li-Cor Biosciences, Lincoln, NE). Briefly, 20 μL of the lysate resuspended in SDS-PAGE sample buffer were boiled for 5 min at 95°C and run at 150 V for 60 minutes on a denaturing non-reducing NuPAGE 4 – 12% Bis-Tris SDS-PAGE gel (Thermo Fisher) and later transferred to PVDF membrane (Bio-Rad, Hercules, CA, USA). The membrane was blocked for 1 hour with Intercept blocking buffer at room temperature and then incubated with Quick Western Detection Solution. This detection solution contains the primary antibody of choice diluted according to the manufacturer, IRDye 680RD detection reagent (1:1000 dilution) in Intercept blocking buffer solution, 0.2% Tween20, and 0.02% SDS. The antibodies used as primary antibodies are: CD36 (Human CD36/SR-B3; Cat no: AF1955) diluted to 1:1000 or CEACAM-1 (Human CEACAM-1/CD66a; cat no: MAB22441) diluted to 1:2000. Both antibodies were purchased from R&D Systems. The membrane blot was incubated overnight at 4°C on a platform shaker. After the incubation, the detection solution was carefully decanted, and the membrane was rinsed 3 times with 15mL of 1X PBS-T (1X PBS + 0.1% Tween 20) and scanned with the Odyssey Imaging System using the 700 nm channel.

### AlphaFold models

We used AlphaFold v3.0 installed on a GPU cluster and obtained 50 models for each system. CD36 and CEACAM1 sequences were obtained from Uniprot. The chemical formula for LPS-Lipid-A was supplied to AlphaFold in the form of a SMILES string.

### Molecular Dynamics simulations of open form of CEACAM1 N domain

The starting structure for the CEACAM1 N domain was derived using AlphaFold v3.0 by submitting the protein sequence to the AlphaFold Server (https://alphafoldserver.com) (42). The input for MD simulation was prepared using the tleap module of AmberTools (43). The protein structure was solvated in a rectangular water box of dimensions 94Å x 81Å x 81Å and Na^+^/Cl^-^ ions were added to neutralize the total charge and achieve an effective salt concentration of 0.15M. The system was parameterized using the FF19SB forcefield and the TIP4PEW water model (44,45). The system was minimized for 2000 cycles of which, first 1000 were via steepest descent and the rest using the conjugate-gradient method. Next, the system was gradually heated from 0 to 310K over 30 ns at constant volume, during which, the protein heavy atoms were restrained by applying a harmonic force with force-constant 500 kcal/mol/Å. Following this, the system was simulated for a further 50 ns in the NPT ensemble (310K, 1 atm) during which, the harmonic restraints were gradually reduced to zero. A final equilibration run was performed for 50 ns without any restraints. The production runs consisted of two independent simulations lasting for 1.8-1.9 µs starting with randomly generated velocities. Temperature and pressure were maintained using the Langevin thermostat with collision frequency gamma ln set to 1.0 The MD simulations were performed using GPU accelerated AMBER v22. Hydrogen mass repartitioning was applied to enable a timestep of 4 femtosecond (46).

### Molecular dynamics simulations of D82A mutant of glycosylated CEACAM1-N domain

In the AlphaFold derived structure of human CEACAM1-N domain, D82 was mutated to alanine by direct deletion of the aspartate side chain and changing the residue name to ALA in the pdb file. The structure of the glycosylated CEACAM1-N domain was derived using the Glycoprotein Builder web interface at GLYCAM-Web (https://glycam.org/gp/)(47), where the positions N77 and N81, the sites nearest to D82, were glycosylated. The MD system was prepared using the Glycan Modeler module of CHARMM-GUI (48). The system was solvated in a water box of dimensions 92 Å X 92 Å X 92 Å and Na^+^/Cl^-^ ions were added to neutralize the net charge and achieve an effective salt concentration of 0.15M. The system was parameterized using the CHARMM36m forcefield (49) and simulated using the GPU accelerated AMBER v22.

Equilibration started with 5000 cycles of minimization of which, the first 2500 were via steepest descent and the rest using the conjugate-gradient method. Following this, the system was heated to 310K over 125 ps and then equilibrated for 10 ns in the NVT ensemble with a timestep of 1 femtosecond. During the minimization and heating phases, the protein heavy atom positions were restrained by applying harmonic forces with force constant 1 kcal/mol. Further, the glycan dihedrals were restrained using harmonic forces with a force constant of 1 kcal/mol. The production run continued for 2.5 μs in the NPT ensemble (310K, 1 atm). Temperature and pressure were maintained using the Langevin thermostat with collision frequency gamma ln set to 1.0 Hydrogen mass repartitioning was applied to enable a timestep of 4 femtosecond (46).

### Modeling of LPS-Ra bound to CEACAM1-N domain

For modeling bound LPS, we selected the MD conformation of the D82A-CEACAM1-N domain showing the largest distance between N81 and N61, ensuring an open conformation with adequate space to dock LPS. Due to the large size and high number of rotatable bonds, the entire LPS molecule could not be docked to CEACAM1 using conventional docking. We therefore adopted a piecemeal approach, where the di-glucosamine segment of lipid-A was first docked to the CEACAM1 cavity using GLIDE SP (50). The ligand conformations were generated outside the protein cavity using MacroModel, followed by the docking of the ligand structural ensemble to CEACAM1. During docking, the vdW radii of both protein and ligand atoms were scaled by 0.5. The best pose with the lowest docking score was selected for the next phase in which the alkyl chains were manually added using the Molecule Builder interface of Maestro, followed by local minimization of the modeled atoms, keeping the protein rigid. Finally, with lipid-A in place, the core glycan moiety was docked to CEACAM-1 and the pose with the lowest docking score was selected. The lipid-A and core glycan were then covalently connected using the molecule builder interface of Maestro. The final structure was subjected to thorough minimization using the PrimeX module of Maestro.

### Location of primary data

Primary data is available from the senior author by request.

## RESULTS

### mLPS-Ra binding to sCEACAM1-4D by SPR

*E coli* mLPS-Ra that contains all of the core glycan and lipid A of LPS was used in these analyses since it retains the endotoxin activity of LPS. mLPS-Ra is a micellar, water soluble LPS, for which an x-ray structure of its monomeric unit in the TLR4-MD2-LPS complex is available (27). To approximate a natural state for mLPS, we made no attempt to disrupt its micellular structure in this study. Since human CEACAM1 is expressed with an IgV-like N-domain followed by 2-3 extracellular IgC domains due to alternative mRNA splicing (1), we expressed a soluble form of the largest splice variant with 4 extracellular domains (sCEACAM1-4D, **Supplemental Fig S1**), preciously described by us (51). When sCEACAM1-4D was immobilized on a CM5 chip and binding studies with LPS-Ra performed over the range of 0-4.0 µM, a K_D_ of 1.04 x 10^-10^ M was obtained (**Fig 1A**). The dissociation of mLPS-Ra from immobilized CEACAM1-4D was very slow, in agreement with the fact that sCEACAM1-4D was binding mLPS-Ra over multiple copies of its subunits resulting in a high avidity interaction. Since the slow dissociation results in an unreliable K_D_ due to rebinding and mass transport effects (52), the binding was also analyzed using a steady state affinity fitting that gave a more reliable value for K_D_ as 1.47 x 10^-6^ M (**Supplementary Fig S2A**).

**Fig 1.**
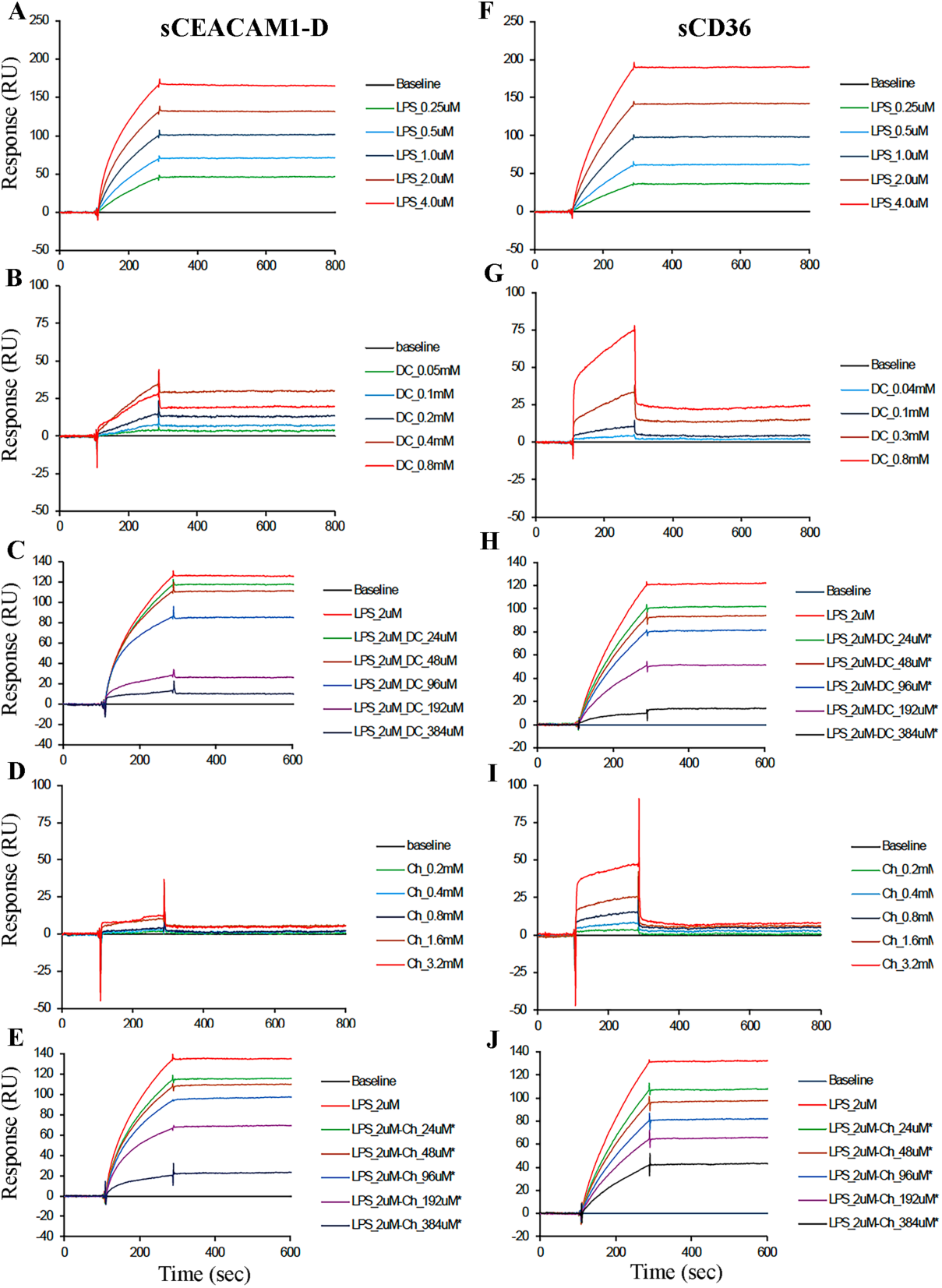
Surface plasmon resonance analysis of the binding of *E. coli* mLPS-Ra or BAs to sCD36 and sCEACAM1-4D. **A.** sCEACAM1-4D (50 µg/ml) was immobilized on a CM5 chip and mLPS-Ra binding performed over the range 0-4 µM. **B-C**. Binding of different concentrations of sodium deoxycholate in the absence (**B**) or presence of mLPS-Ra (**C**) to immobilized sCEACAM1-4D. **D-E**. Binding of different concentrations of sodium cholate in the absence (**D**) or presence of mLPS-Ra (**E**) to immobilized sCEACAM1-4D. **F**. sCD36 (50 µg/ml) was immobilized to a BIACore CM5 chip and binding studies with mLPS-Ra (0 to 4 µM) was performed. **G-H**. Increasing concentrations of sodium deoxycholate in the absence (**G**) or presence of mLPS-Ra (**H**) to immobilized sCD36. **I-J**. Increasing concentrations of sodium cholate in the absence (**I**) or or presence of mLPS-Ra (**J**) to immobilized sCD36. The kinetic constants for LPS binding to sCEACAM1-4D are k_a_ = 5.85x10^3^ 1/Ms, k_d_ = 6.08x10^-7^ 1/s, and K_D_ = 1.04x10^-10^ M. The kinetic constants for LPS binding to sCD36 are k_a_ = 3.69x10^3^ 1/Ms, k_d_ = 1.25x10^-6^ 1/s, and K_D_ = 3.38x10^-10^ M. Each concentration of mLPS-Ra or BA was performed in duplicate (ave. shown).

Since CEACAM1-4D has 4 domains, its interaction with LPS could be with any one or a combination of its domains. Given that the N-domain of CEACAM1 has been shown to convey its most conserved feature, we attempted to block mLPS-Ra binding to the N-domain with an N-domain specific antibody, T84.1 (53). When immobilized sCEACAM1-4D was first subjected to binding T84.1 followed by mLPS-Ra (0.5 µM), the RUs bound were similar to binding of LPS (0.5 µM) alone (**Supplemental Fig S3A**), demonstrating that this antibody did not block LPS binding. Since this experiment did not definitely rule out the N-domain as an the only mLPS-Ra binding domain, we separately expressed CEACAM1-N domain (**Supplemental Fig S3B**) and found mLPS-Ra bound with K_D_ of 3.56 x 10^-9^ M using kinetic fitting model (**Supplemental Fig S3C**) and a K_D_ of 1.40 x 10^-6^ M from a steady state affinity binding approach (**Supplemental Figure S2B**). We conclude that the N-domain is the minimum domain required for mLPS-Ra binding.

### Effect of bile acids on mLPS-Ra binding to sCEACAM1-4D by SPR

To approximate physiological conditions for mLPS binding to CEACAM1, we note that CEACAM1 is expressed in the liver and intestine where it is exposed to bile acids (BAs), and that BAs are known to reduce the micelle size of mLPS (54). These studies were performed with two representative BAs, sodium cholate, a primary BA, and sodium deoxycholate, a secondary BA, in the absence or presence of mLPS-Ra **(Fig 1B-E).** Sodium deoxycholate (0.05 - 0.8 mM) in the absence of mLPS-Ra showed detectable binding to CEACAM1-4D compared to sodium cholate (0.02 - 3.2 mM). In both cases the bound BAs exhibited slow rates of dissociation; however, no reliable rate constants were obtained for these two BAs binding to CEACAM1-4D.

Having established that sCEACAM1-4D binds both mLPS-Ra and BAs, we determined the effect of BAs on a fixed concentration of mLPS-Ra (2 µM) binding to sCEACAM1-4D, with increasing concentrations of deoxycholate (**Fig 1C**). The results indicate that the RU values from mLPS-Ra-deoxycholate mixtures are much less than the sum of RU from individual mLPS-Ra and deoxycholate alone. This could be due to both mLPS-Ra and deoxycholate binding to the same site on CEACAM1-4D, but since the molecular size of deoxycholate is smaller, the binding of deoxycholate instead of LPS-Ra would reduce the RU signal significantly. Another possible explanation is that deoxycholate may reduce the micelle size of mLPS-Ra leading to a reduced RU signal. Using equivalent concentrations of cholate, the study was repeated (**Fig 1E**), indicating that the effect of cholate on LPS binding was less than that of deoxycholate. Since deoxycholate is known to reduce the size of LPS micelles (54), changes in the total binding (measured in RU) of mLPS-Ra to CEACAM1-4D in the presence of BAs is a reasonable explanation.

### mLPS-Ra binding to sCD36 by SPR

Since our previous studies showed that CEACAM1 and CD36 co-associate in hepatocytes (23) and CD36 was known to affect LPS uptake in the intestine (22), we asked if we could demonstrate direct binding of mLPS-Ra to sCD36, similar to sCEACAM1-4D. When soluble sCD36, obtained from a commercial source that was previously validated by SPR binding studies to oxLDL (55), was immobilized on a CM5 chip and binding studies with mLPS-Ra performed over the range of 0-4.0 µM, a K_D_ of 3.38 x 10^-10^ M was obtained (**Fig 1F**). Although the kinetic association constant k_a_ for sCD36 was similar to that for sCEACAM1-4D, the dissociation constant k_d_ for both sCD36 and sCEACAM1-4D were too small to be determined precisely, indicating both were binding micelles with large mass transport effects. When the K_D_ for mLPS-Ra binding to sCD36 was recalculated using a steady state affinity binding approach, a value of 2.23 x 10^-6^ M was obtained (**Supplemental Fig S2C**), slightly larger than that for sCEACAM1-4D or N-domain.

### Effect of bile acids on mLPS-Ra binding to sCD36 by SPR

Similar to CEACAM1-4D, CD36 is highly expressed in the intestine (22) and exposed to BAs during digestion. When BA binding studies were performed with immobilized sCD36 in the presence or absence of mLPS-Ra, we found in the absence of mLPS-Ra, higher binding response of both deoxycholate and cholate to sCD36 than that to CEACAM1-4D, while the effect of both bile acids on the mLPS-Ra binding to sCD36 share the similar pattern with binding to CEACAM1-4D **(Fig 1G-J).** Since it appears that BAs compete for the same binding site on sCD36, or CEACAM1-4D we replotted the data as normalized RU (± correction for the binding of BA only) vs the molar ratio of BA/mLPS-Ra. In the case of sCD36, a smooth curve was obtained for cholate, indicating that cholate reduced the mLPS-Ra binding RUs mainly by one mechanism, that is reduction of the micelle size of mLPS-Ra. For the curve of deoxycholate there is a discontinuity around a molar ratio of 50, suggesting that in addition to the reduced size of mLPS-Ra micelles, the deoxycholate also competes with the same binding site on sCD36 as its concentration increases. (**Supplementary Fig S4A**). Since deoxycholate is more efficient than cholate in reducing the size of mLPS micelles (54) these results are expected. Similar results were obtained for the effect of BAs on sCEACAM1-4D with cholate giving essential linear results with deoxycholate giving higher order curves (**Supplementary Fig S4B**). In both cases, deoxycholate was more effective in suppressing mLPS-Ra binding to both proteins.

### Binding of mLPS-Ra to sCD36 or sCEACAM1-4D by SEC

To determine the binding of mLPS-Ra in solution phase, sCD36 (35 µg/0.1 mL; 5 µM) or sCEACAM1-4D (35 µg/0.1 mL; 2.9 µM), were incubated with increasing amounts of mLPS-Ra (0-150 µg/0.1 mL) for 60 min at 37°C, then the entire volume (0.1 mL) analyzed by SEC. It should be noted that mLPS-Ra is UV transparent and thus only changes in the retention times of sCD36 or sCECAM1-4D were observed. The results show that both sCD36 and sCEACAM1-4D generate higher molecular complexes with mLPS-Ra that reach a maximum at about 100 µg of mLPS-Ra (**Fig 2**). Since the injected complex undergoes a dynamic dilution of about 10-fold during chromatography, it is possible that a certain amount of complex dissociation occurs preventing conversion to 100% complex formation. In any case, the molecular size of the CD36-mLPS-Ra complex was estimated by reference to standards of known Stokes radii as 7.9 nm (ca 500 kDa for a globular particle) and 8.9 nm (800 kDa) for the CEACAM1-4D-LPS-Ra complex (**Supplemental Fig S5**). Whereas sCD36 can be approximated as a globular structure, that is not the case for sCEACAM1-4D that was shown by us to be a prolate ellipsoid with an axial ratio of 8 and a Stokes radius of 5.5 nm by ultracentrifugation (51). In good agreement, the calculated Stokes radius of CEACAM1-4D from **Supplemental Fig S5** is also 5.5 nm.

**Fig 2.**
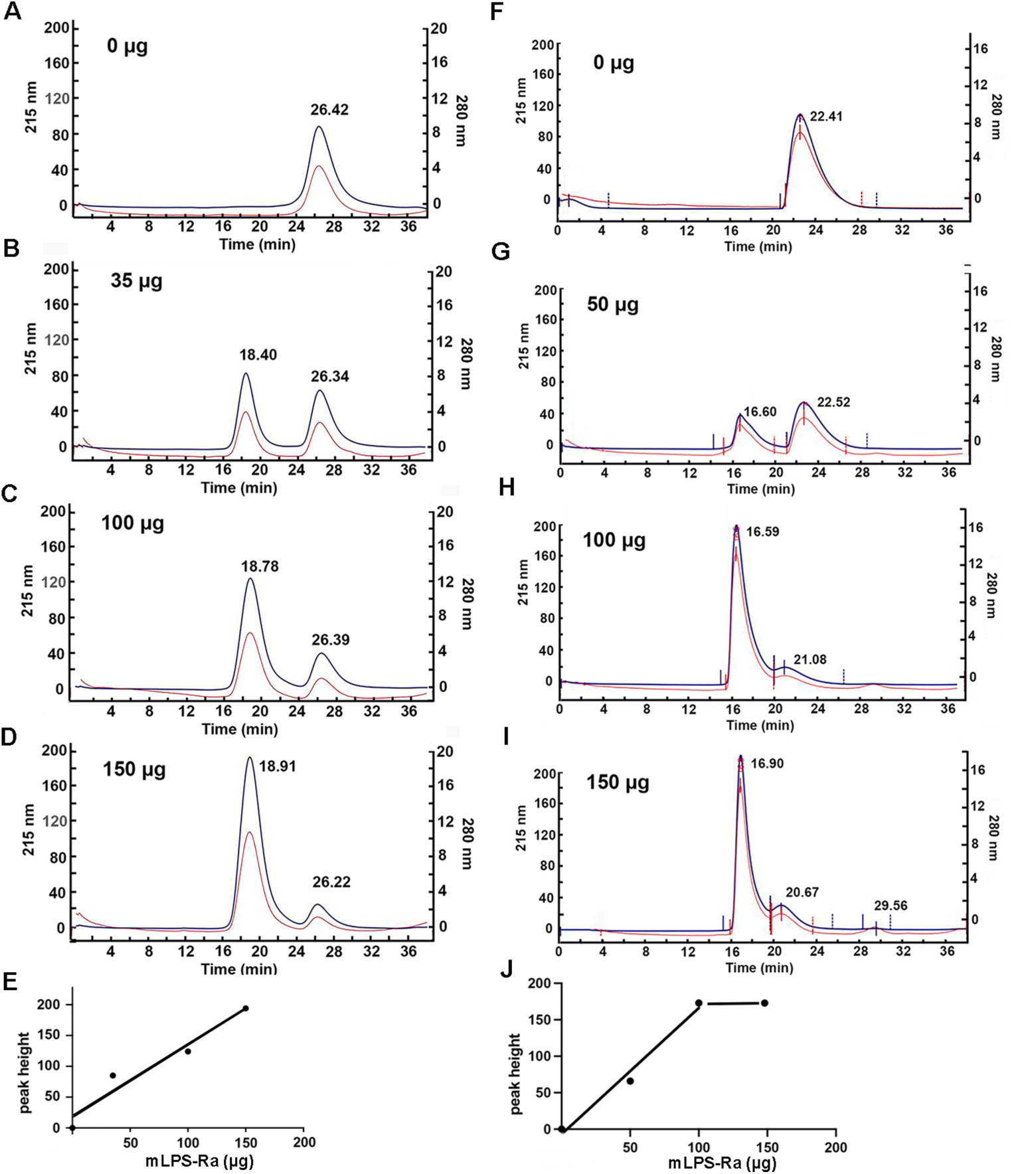
Size exclusion chromatography analysis of the binding of *E. coli* mLPS-Ra to sCD36 and sCEACAM1-4D. **A-D.** sCD36 (35 µg, 0.5 nmol) was incubated with increasing amounts of mLPS-Ra (0-150 µg, 0-30 nmoles assuming a monomeric molecular weight of 5 kDa) for 1 hr at RT in a volume of 0.1 mL and the entire amount injected onto a Superdex 200i column in PBS at a flow rate of 0.5 mL/min monitored at 215 and 280 nm. **F-I**. sCEACAM1-4D (35 µg, 0.29 nmoles) was incubated with increasing amounts of mLPS-Ra and analyzed as described in **A-D**. Note: mLPS-Ra is UV transparent at 215 nm in these studies. **E** and **J** plots of the peak heights of sCD36 and sCEACAM1-4D vs the concentration of mLPS-Ra.

### Particle sizes of LPS-Ra complexes with sCD36 or sCEACAM1-4D by TEM

Direct visualization of sCD36-LPS-Ra complex from SEC (see above) was performed by TEM: a range of ellipsoid particle sizes were observed: with a median minor axis of 8.4 nm and major axis of 12.3 nm (**Fig 3A**). The calculated radius (6.1 nm) of the median particle is less than the Stokes radius obtained by SEC (7.9 nm), while the maximum radius of 8.6 nm is larger than obtained by SEC. However, the gaussian distribution of particle sizes seen in TEM agrees well with the symmetrical peak seen on SEC. Similar results were observed for the mLPS-Ra-CEACAM1-4D complex: a median minor axis of 7.4 nm and major axis of 12.0 nm (**Fig 3B**). While the median radius calculated from TEM (6.0 nm) does not agree with that obtained from SEC (8.9 nm), the maximum radius (8.9 nm) is in good agreement.

**Fig 3.**
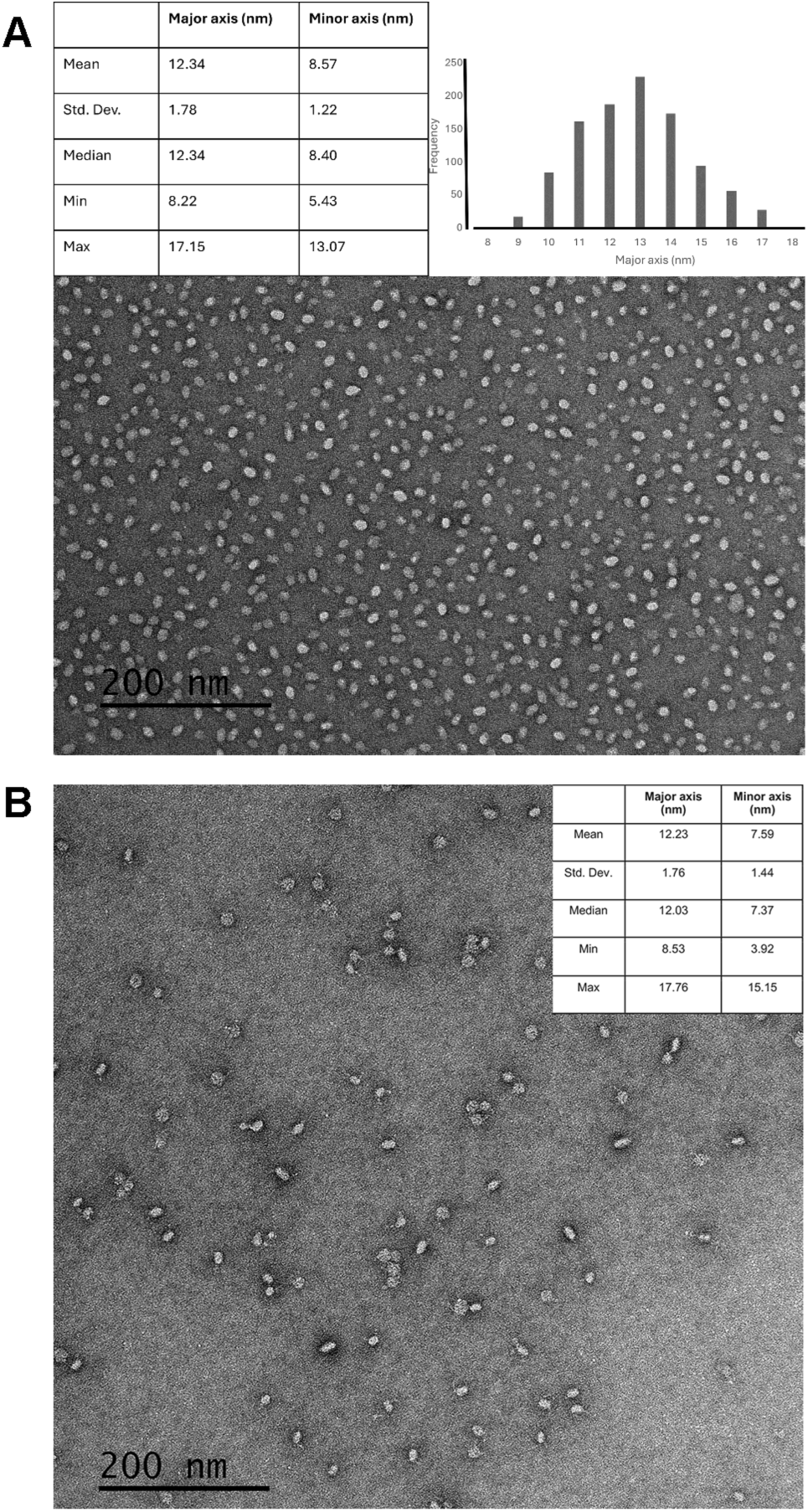
Transmission electron microscopy analysis of *E. coli* mLPS-Ra complexes with sCD36. **A-B.** The sCD36-mLPS-Ra (**A**) and sCEACAM1-4D-mLPS (**B**) complexes isolated from SEC (see Fig 2) were analyzed by TEM. Particle size (n> 200) analysis shows measurement and distribution of ellipsoid complexes.

### Interaction of sCEACAM1 with sCD36 by SPR

CD36 and CEACAM1 are co-expressed in several cell types that respond to LPS, including hepatocytes (56,57) and macrophages (58,59). In the case of intestinal enterocytes, the primary source of nutritional fatty acid uptake, both CEACAM1 and CD36 are expressed (20,60) and CD36 has been shown to uptake LPS along with fatty acids (20). The possibility that the two receptors could associate with each other was established by us in HepG2 hepatocytes by co-immunoprecipitation with specific antibodies (23). Indeed, SPR studies showed direct binding of immobilized sCD36 with sCEACAM1-D with a K_D_ of 5.28 x 10^-8^ M. (**Fig 4A**). However, when the two soluble proteins were incubated together and run on SEC, there was no evidence of complex formation. This is not a surprising result given that CEACAM1 dimer formation requires it to be in the membrane with a prior interaction between its TM domains (4). As a first approach, we used Alpha Fold to dock the two proteins and then placed them in a model membrane. Among an ensemble of docked structures (not shown), only one was compatible with bringing their TM domains in close proximity (**Fig 4B**). In order to test this model, we performed two types of proximity labeling assays (PLA).

**Fig 4.**
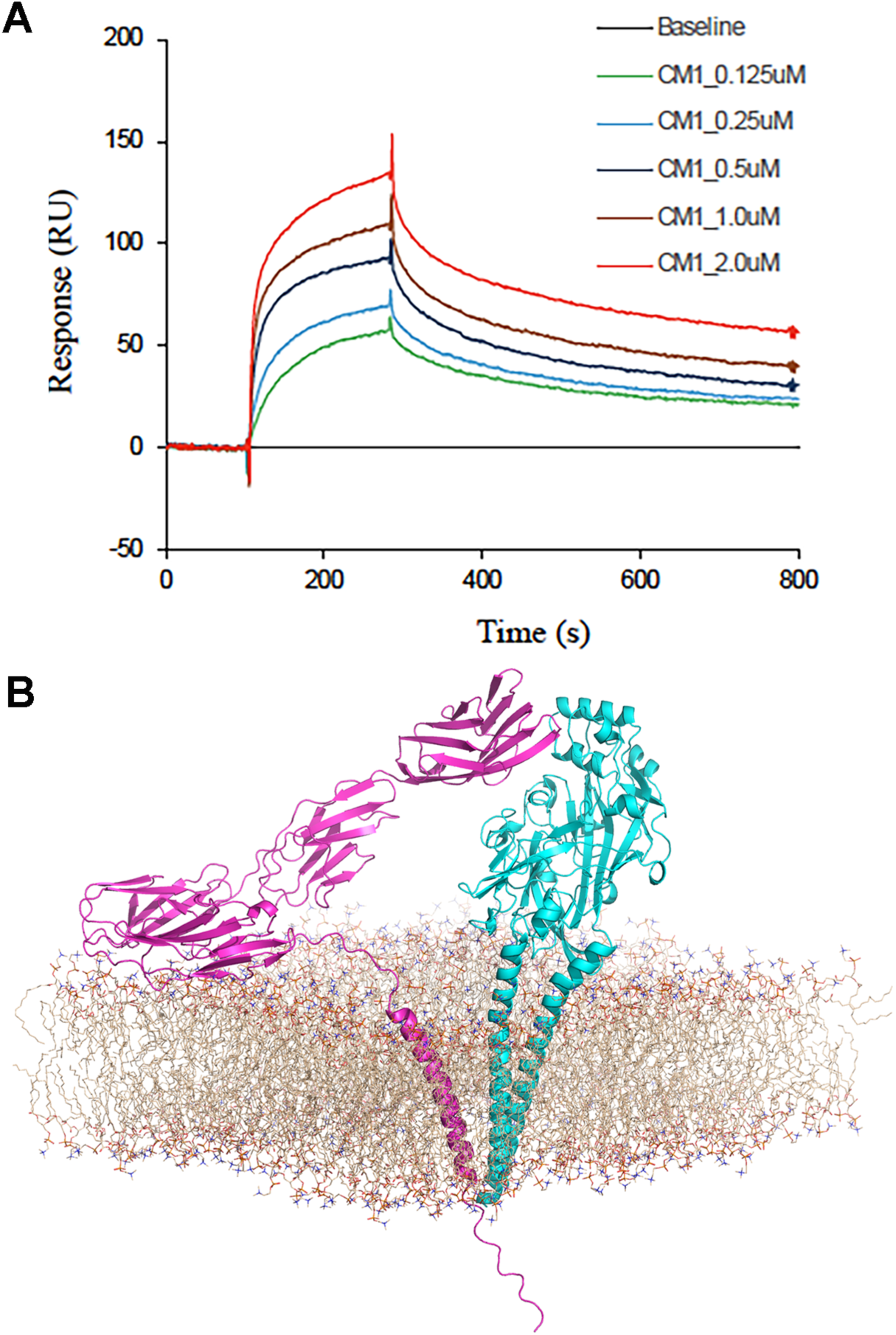
Direct binding of sCEACAM1-4D to sCD36 and Alpha Fold model of CEACAM1-4D cis interaction with CD36. **A.** sCD36 (12.5 µg/ml) was immobilized to a BIACore CM5 chip and SPR binding studies with CEACAM1 (0-2.0 µM) performed. Chips were regenerated between runs with 0.1 N HCl. The kinetic constants are k_a_= 3.25 x 10^4^ M^-1^s^-1^ and k_d_= 1.72 x 10^-3^ s^-1^ with K_D_ = 5.28 x 10^-8^ M. **B.** Alpha fold model of interaction of CEACAM1-4D (magenta) with CD36 (cyan) in a model membrane (brown).

### Proximity ligation of CEACAM1-4L with CD36

In the first type of PLA, specific antibodies to CEACAM1 and CD36 were ligated with complementary oligonucleotides that served as a template for amplification with fluorescent labeled probes on HepG2 hepatocytes that expressed both CEACAM1-4L (CEACAM1 with four extracellular domains and a 72-amino acid long cytoplasmic domain) and CD36 or CD36 only as a control. Analysis of the cells by confocal microscopy after PLA showed a strong cell surface signal (P< 0.001) for the CEACAM1-CD36 double positive sample vs the negative control lacking CEACAM1 (**Fig 5A-C**). In addition, HEK cells double transfected with plasmids for full length CEACAM1-4L and CD36 showed strong signals (P< 0.01) in the PLA assay vs untransfected HEK controls (**Fig 5D-F**). Flow analysis of the HEK cells for none, single, and double transfections are shown below. We conclude that the ectodomains of both proteins can associate with each other at the cell surface.

**Fig 5.**
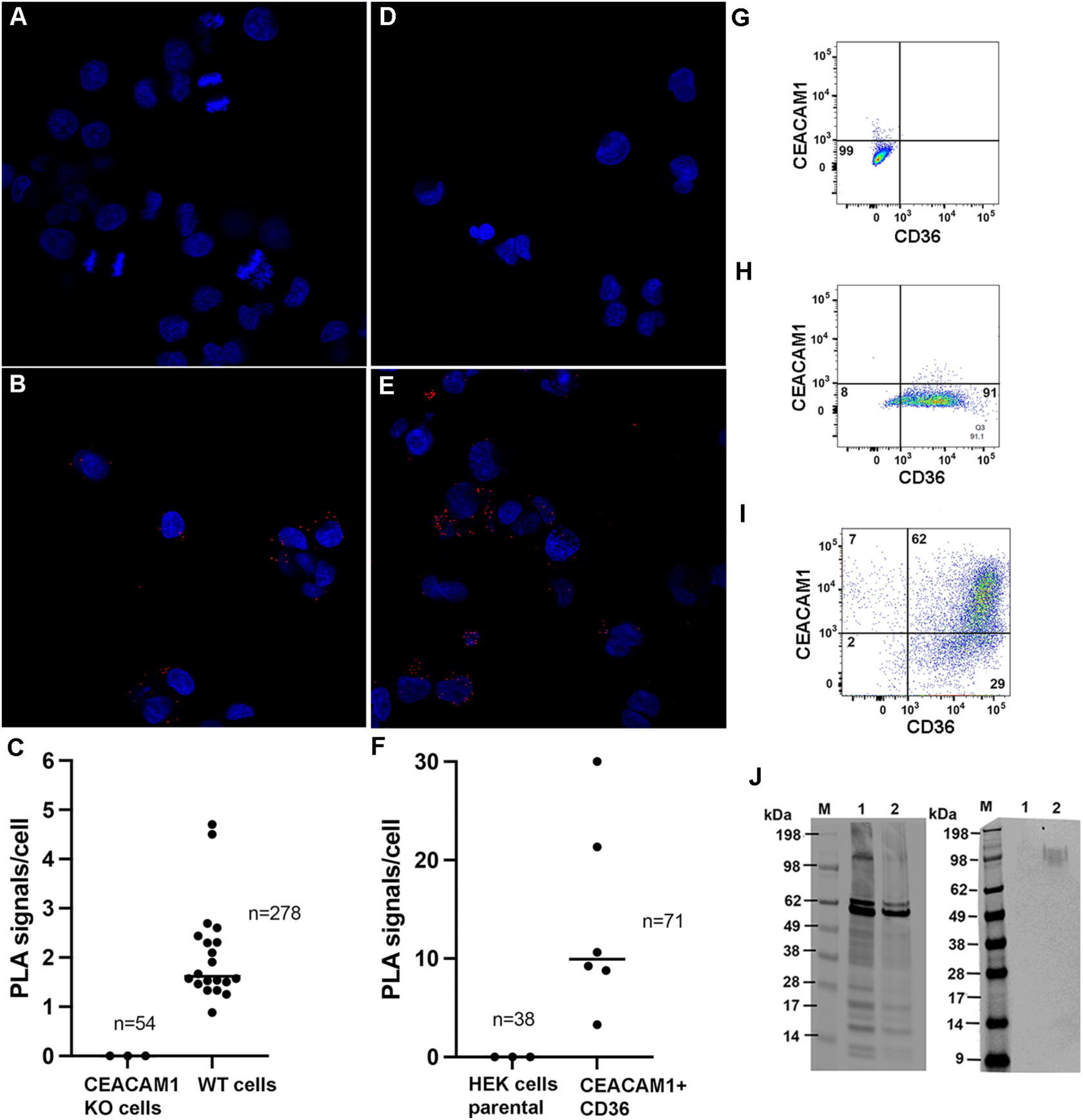
Proximity ligation assays for CD36 and CEACAM1-4L. **A-B.** HepG2 hepatocytes expressing only CD36 (**A**) or both CD36 and CEACAM1-4L (**B**) were cross-linked with CD36 and CEACAM1 specific antibodies conjugated with complementary oligonucleotides and amplified with red fluorescent labeled primers. **C**. Quantitation of PLA signals for **A** (n=54) and **B** (n=278) with a P< 0.0001. (**D-F)**. HEK cells expressing only CEACAM1-4L (**D**) or both CEACAM1-4L and CD36-BioID2 (**E**) were treated as in **A-B**. **F**. Quantitation of PLA signals for D (n= 38) and E (n= 71) with a P< 0.018. Cells were counterstained with DAPI (blue). **G-J.** Flow analysis of CEACAM1-4L and CD36-BioID2 expressions in transfected HEK cells. (**G**) untransfected controls. (**H**) HEK cells transfected with CD36-BioID2 only. (**I)** HEK cells transfected with both CD36-BioID2 and CEACAM1-4L. **J**. Immunoblots of HEK cells expressing only CD36-BioID2 (lane 1) or both CD36-BioID2 and CEACAM1-4L (lane 2), immunostained with antibodies to CD36 (left) or CEACAM1-4L (right).

In a second PLA assay involving biotinylation of adjacent cytoplasmic domains by BioID2 (61), HEK cells were transfected with plasmids expressing a C-terminal CD36-BioID2 fusion protein only, or CD36-BioID2 fusion protein (**Supplemental Fig S6**) plus CEACAM1-4L (37) followed by flow analysis with CEACAM1 and CD36 specific antibodies (**Fig 5G-I**). Since the flow analysis revealed a good co-expression of the double transfectants, we proceeded with the proximity labeling assay. The cell double transfectants were incubated with biotin, lysed, the biotinylated proteins isolated on streptavidin beads. The bound and eluted proteins were immunoblotted with anti-CD36 or anti-CEACAM1 antibodies (**Fig 5J**). The results show that the CD36-BioID2 transfectants are biotinylated in the singly transfected cells, and both CEACAM1 and CD36-BioID2 co-transfectants are biotinylated in the double transfected cells. We conclude that the cytoplasmic domains of both proteins associate with each other.

The fact that two different PLAs, one directed at the extracellular side of the plasma membrane, and one directed at the cytoplasmic side of the plasma membrane, show CD36 and CEACAM1 are expressed in close proximity to each other, suggests that they can cooperate in cell signaling. This may explain their similar roles in regulating LPS-TLR4 signaling and their ability to bind mLPS-Ra micelles. In addition, these results provide evidence for a cis-model of interaction (**Fig 4B**). The possibility that a trans-interaction between cells may also be possible requires further studies.

## DISCUSSION

While it was known that bacterial adhesins bind CEACAM 1 (62), it was first shown by Servin and co-workers (10) that CEACAM1 transfected epithelial cells bind LPS and that bacterial expressed CEACAM1 N-domain bound LPS. Despite this proposed novel function of CEACAM1, the finding received little attention, perhaps due to the lack of a structural basis for this interaction. In this regard, we noticed that MD2, that was crystalized with the LPS analog lipid IVa, enveloped lipid IVa within the hydrophobic interior of its beta-sheet sandwich (30), a feature it shares with the N-domain of CEACAM1. It is noteworthy than both MD2 and the N-domain of CEACAM1 lack an internal disulfide bond that covalently links the beta sandwich in most IgV-like analogs, and if present, would prevent the accommodation of lipids within their hydrophobic interiors. However, a unique feature of CEACAM1’s N-domain is the presence of a salt bridge between residues Asp-82 and Arg-64 (63) that we propose must be disrupted to accommodate its interaction with mLPS. To simulate such a disruption, we performed molecular dynamic analysis on a Asp82Ala N-domain mutant that predicted a stable formation of an open conformation that accommodates LPS-Ra within its beta-sheet structure (**Fig 6**). In order to explain why CEACAM1 is still attached to mLPS (according to our SPR, SEC and TEM studies), the model needs to be expanded with a structural representation of the LPS micelle.

**Fig 6.**
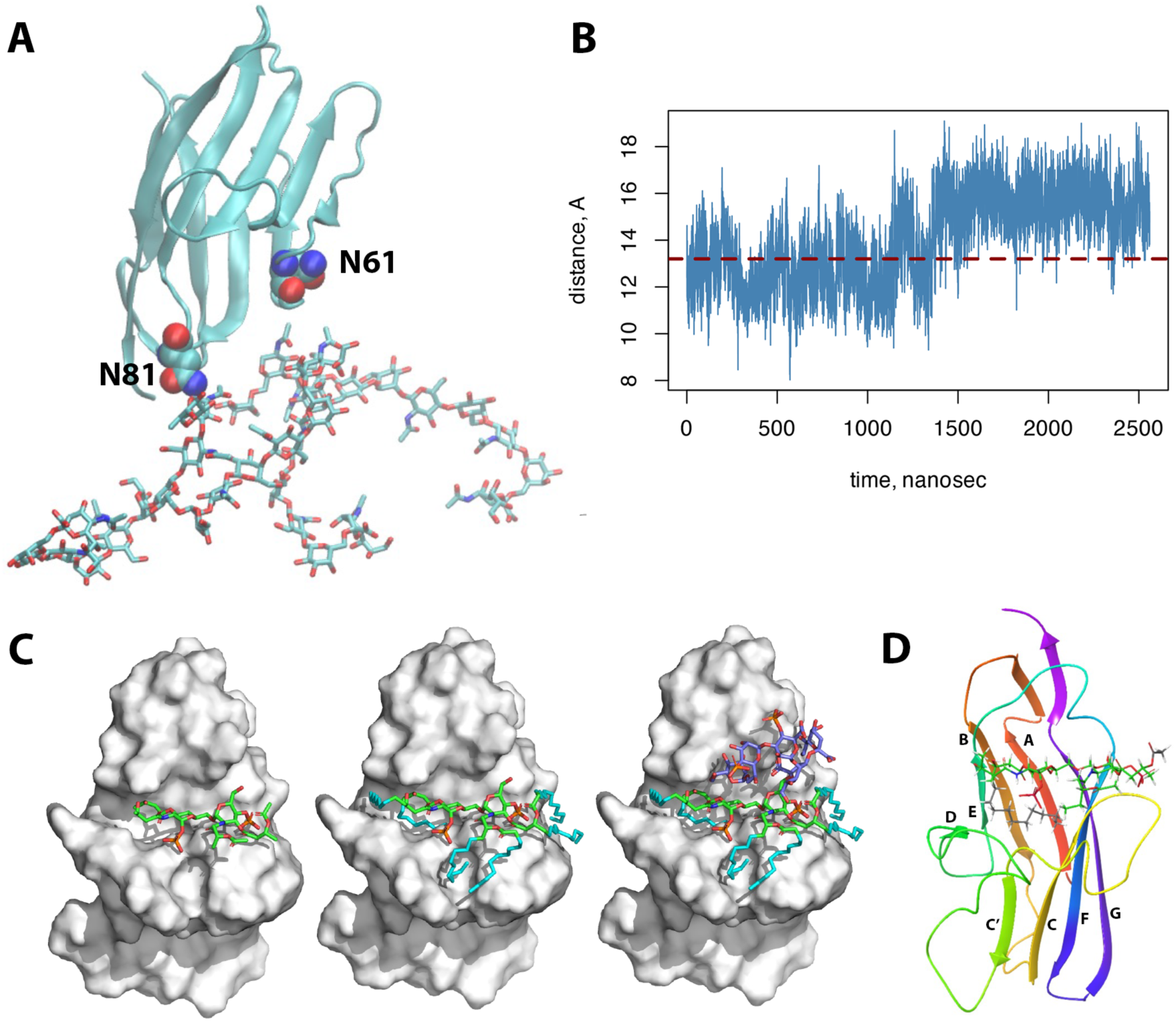
Molecular dynamics model of D82A mutant of CEACAM1-N domain with a minimum LPS-Ra unit. **A**. Alpha Fold model of D82A mutant of CEACAM1-N showing distance markers residue N81 with attached glycan and N61. **B.** MD simulation of distance between two marker residues over 2 x 1 µsec simulation. **C.** Using the open form (greatest distance between markers) from **B**, LPS-Ra unit was sequentially (left to right) docked into a surface model of CEACAM1-N. **D.** Ribbon model with beta-strands shown (A to G) of CEACAM1-N with docked structure of LPS-Ra unit.

Generating a model of mLPS binding to CD36 was less of a challenge, since CD36 has a hydrophobic pocket that has already been shown to accommodate long chain fatty acids such as palmitic acid (64). Using the newest version of Alpha Fold we were able to dock two acyl chains of the Lipid A portion of LPS-Ra into the hydrophobic pocket of CD36 (**Fig 7**). Since the model predicts that the additional four acyl chains of Lipid A are unbound, we predict that they remain attached to the mLPS. Such a model would explain our binding studies that show association of CD36 with mLPS.

**Fig 7.**
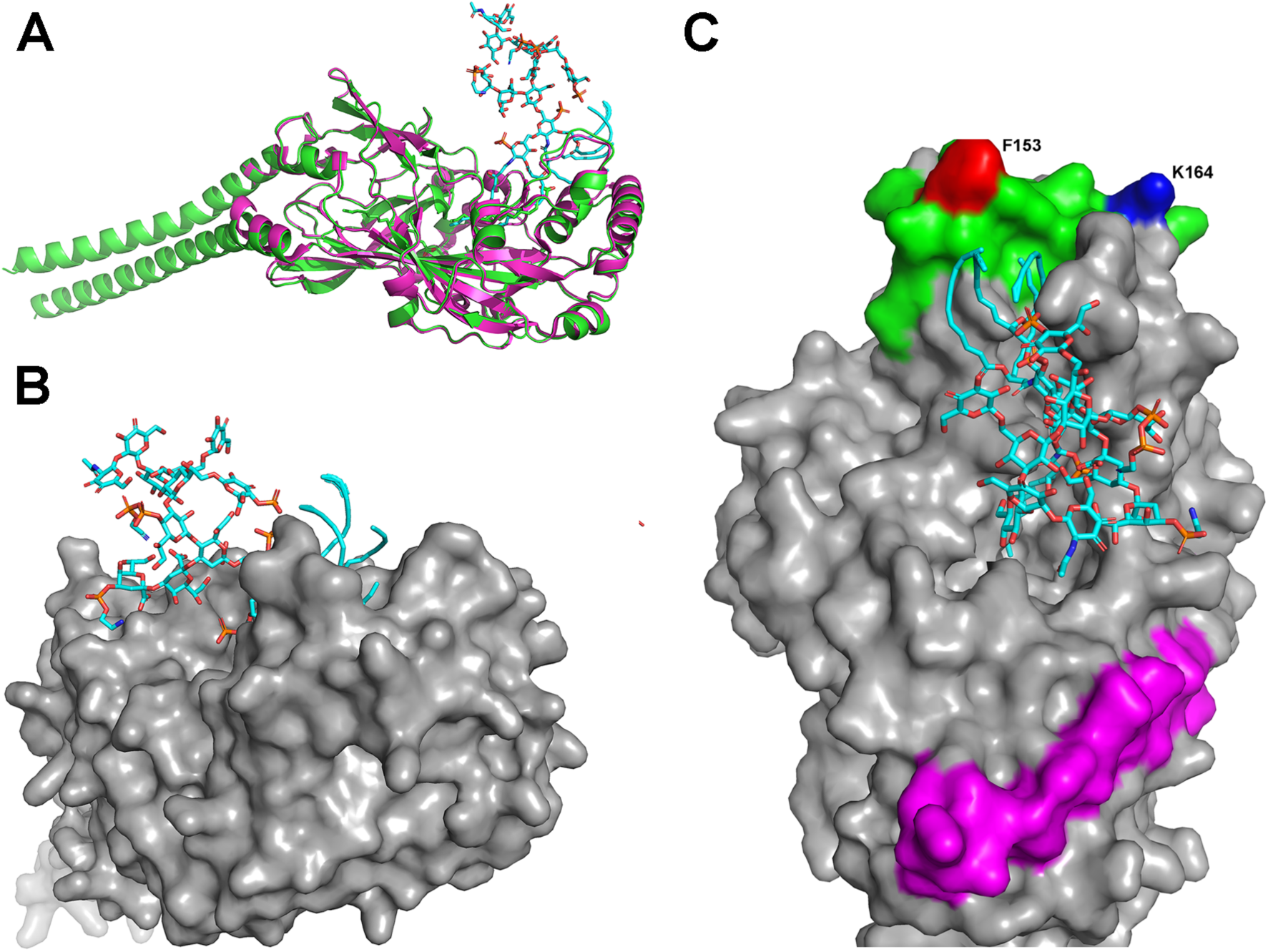
Model of LPS-Ra binding to CD36. **A**. Ribbon model of CD36 with stick model of LPS-Ra docked to CD36 (green: full length CD36, Alpha Fold model; magenta: sCD36 x-ray structure). **B**. Surface model view showing LPS glycan and four acyl chains of one of the glucosamine subunits of Lipid A projecting away from the protein surface. **C**. Surface model view showing projection of two acyl chains of one of the glucosamine subunits of lipid A into the protein interior, along with K164 (blue), the long chain fatty acid binding site of CD36 (green) and the binding site of TSP-1 (magenta). The position of F153 (red), the key binding site to the malaria protein is shown as a reference.

The size and milleu of mLPS is an important consideration in understanding the mechanism of mLPS binding proteins such as CEACAM1 and CD36. In the case of the intestine, as well as bile canaliculi, mLPS is likely to interact with bile acids. When studying the effect of bile acids on LPS-Ra binding to either CEACAM1 or CD36, we found that deoxycholate was more effective than cholic acid in reducing the micelle size, a result that agrees with older studies using sedimentation equilibrium (54). A simple 2D model suggests that four bile acids bind the minimum micelle of LPS-Ra (**Fig 8A**). Building on this model, we envision that abundant large micelles are slowly broken up with the BAs at hydrophobic acyl exposed ends, leading to a slow accumulation of the minimum micelle model. Thus, when CEACAM1 or CD36 encounter BA-mLPS complexes, they may replace the BA at the ends of the micelle. In our SPR studies with mLPS plus BAs, we observe less mass (measured in RUs) binding to these immobilized proteins compared to mLPS in the absence of BAs. In the studies without BAs, we predict that CEACAM1 or CD36 bind to the micelle ends displacing any loosely bound LPS monomers that otherwise cap the micelle ends. Although we cannot discern where CEACAM1 or CD36 are located within the complexes on TEM, the shape of the complexes suggests the micelles consist of multiple layers forming well-defined ellipsoid structures.

**Fig 8.**
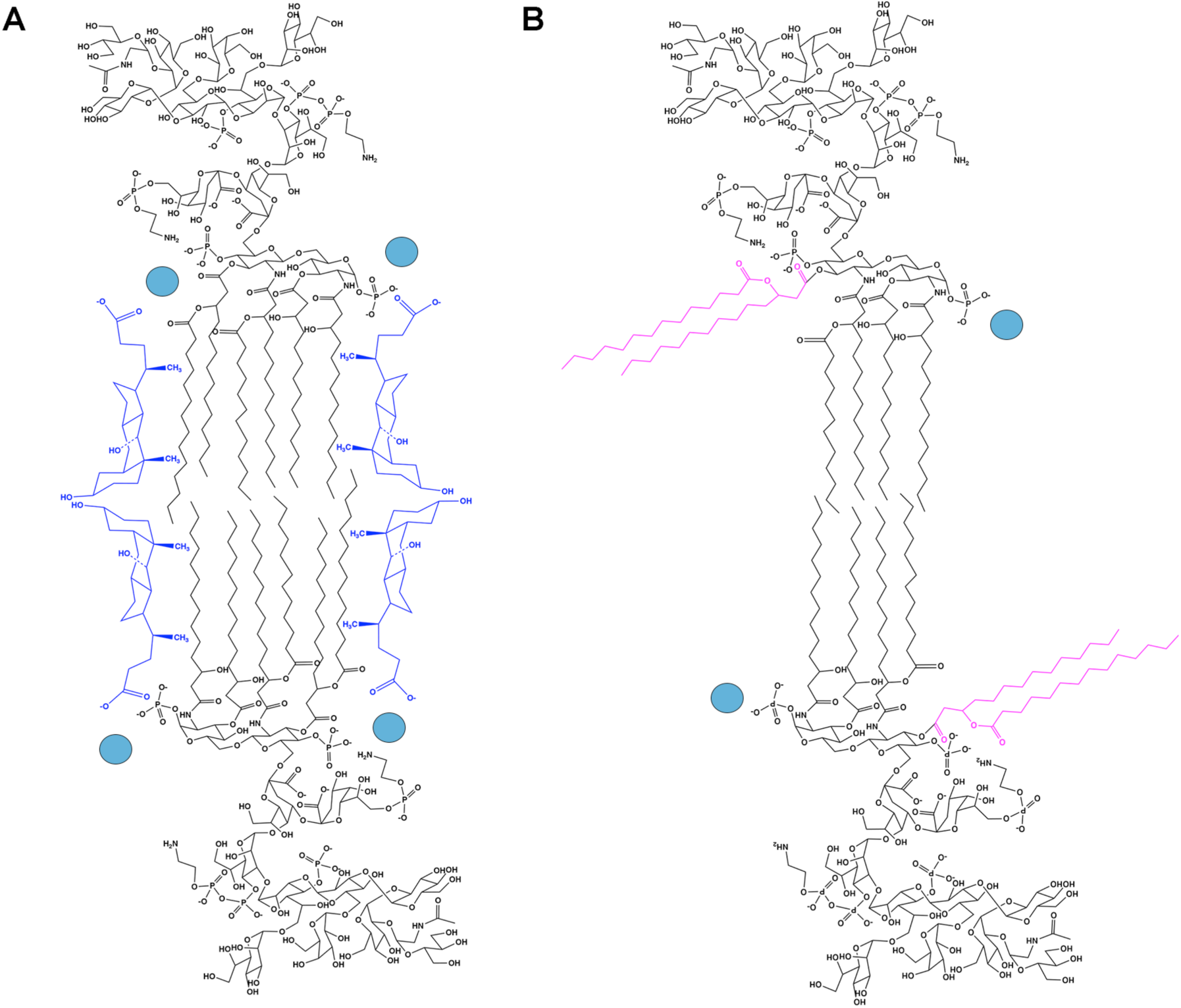
Model of minimum LPS-Ra micelle with and without deoxycholate. **A.** Model of minimum LPS-Ra units with exposed hydrophobic acyl chains capped with deoxycholic acid (dark blue) and calcium ions (light blue). For clarity, two additional molecules of LPS-Ra in ±z-axis directions not shown). **B.** Model of minimum LPS Ra micelle showing rotation about glycosidic bond between hexosamine units exposing two acyl chains (magenta) that may insert into the hydrophobic core of CD36 (CD36 not shown). Calcium ions bridging phosphates shown in light blue.

If we generalize that LPS binding proteins start their interaction with mLPS at its ends where the micelle structure is vulnerable to attack, we propose that a portion of the core structure of Lipid A may be flexible allowing proteins to bind just the portion of Lipid A containing the two acyl chains (**Fig 8B)**. This may explain a unique feature of Lipid A in LPS-Ra in which each repeating unit has two glucosamine residues, one with two acyl chains and the other with four acyl chains. Assuming rotation about the glycosidic bond linking the two glucosamine residues, two of the acyl chains would be available for interactions with LPS binding proteins at the ends of the micelle, while the second glucosamine residue with four acyl chains would remain in the micelle. If this is accurate, then mechanistically, it may explain how LPS binding proteins bind at mLPS ends and “nibble away” at the micelle reducing its size. What makes CEACAM1 and CD36 unique is that they remain attached to the micelle ends.

The expansion of the *CEACAM* gene family across numerous species (7) is evidence of a common function such as mLPS binding that later evolved to include immune signaling to control inflammation. This function may also explain why many strains of pathogenic bacteria have developed adhesin proteins that bind CEACAMs (62,65), allowing them to subvert their host mLPS detection/protective functions, and at the same time, invade the host epithelial barrier. Since the presence of systemic mLPS is associated with life-threatening sepsis (66), it is not surprising that T- and B-cells, as well as granulocytes and macrophages, evolved to express both TLR4 and CEACAM1 to restrain runaway inflammation (14). Thus, even though both TLR4 and CEACAM1 detect the same ligand, namely mLPS or its “bite size” ends, CEACAM1 inhibits the downstream signaling of TLR4, not the actual binding step. CEACAM1 does this by expression of the ITIM containing long cytoplasmic isoform that is missing in epithelial cells that predominantly express the short cytoplasmic domain (67,68).

The fact that the structurally distinct CEACAM1 and CD36 share in binding mLPS and can interact with each other is reminiscent of the role of multiple LPS binding proteins in the transfer of LPS monomers to TLR4. This common theme suggests that mLPS *per se* is not dangerous until it exceeds a threshold, otherwise the host would be in a constant state of inflammation. Indeed, abrogation of either the *Ceacam1* or *Cd36* gene in mice leads to smoldering inflammation that increases with age and lipid consumption where mLPS can masquerade as a hitchhiker. In the case of liver specific *Ceacam1* LKO mice (69), just as in global *Ceacam1* KO mice (70), its absence leads to obesity, insulin resistance and kidney and vascular pathologies *even on a normal diet over time*. While the functional absence of CD36 in certain human populations does not cause life threatening pathologies (71), there are no examples of loss of the human *CEACAM1* gene, suggesting that its function is essential to life. Since mice have two alleles of *Ceacam1*, as well as a second gene (*Ceacam2*), the loss of one is compensated by the expression of the other (72). It is also worth mentioning that LPS species from many commensal bacteria are unable to deliver an inflammatory response via TLR4 signaling, suggesting that the engagement of different species of LPS with CEACAM1 and CD36 may be worth studying. Furthermore, the expression of both CEACAM1 (73) and CD36 (74) on the surface of platelets is functionally important, linking mLPS to capillary clotting during infection (75,76).

Although we focused our studies on the binding of mLPS-Ra to CEACAM1 and CD36, we hypothesize that their binding functions are not limited to mLPS from gram negative bacteria, but could also extend to gram positive micellar lipids such as teichoic acid and TLR1/2 binding lipopeptides such as Pam3CSK4. In this respect, CEACAM1 is known to associate with TLR2 (77), supporting this speculation. Therefore, further lipid binding studies with CEACAM1 are required, as well as their complexes with bile acids. In addition, we speculate that each of the bile acids may have a preferential binding to each of the bacterial lipids, partly explaining their species specificity in different organisms, reflecting not only their diets but also their diet’s contamination with different bacteria. In this respect, we observed a larger effect of deoxycholate on the LPS-Ra micelles binding to sCEACAM1 and sCD36 than that of cholate. Although deoxycholate is a bacterially produced secondary bile acid, its concentration in bile is nearly as high as cholate itself (78). The significance of this observation may indicate either an evolutionary symbiotic relationship or an arms race between bacteria and man.

Finally, it is worth pointing out that chronic over consumption of nutritional lipids that can lead to obesity and type 2 diabetes (T2D) is associated with high levels of circulating LPS (79,80). Moreover, the reduced CEACAM1 expression in the liver of obese patients (70) may contribute to elevated levels of LPS. Interventions such as gastric sleeve reduction of stomach volume has been shown to not only reverse T2D in mice (81) and in man (82), but also can return serum LPS levels to normal (83). This apparent causal relationship among LPS, T2D and gastric sleeve surgery suggests that management of LPS uptake/excretion in the intestine and liver may be an excellent target for the prevention and management of T2D.

## CONCLUSION

The in vitro demonstration of the direct binding of mLPS to CEACAM1 and CD36 by three approaches provides evidence that mLPS binding may operate as an important first step in the management of this ubiquitous bacterial agent. The fact that CEACAM1 and CD36 associate with each other calls attention to a cooperative, physiological relevant mechanism that involves mLPS, and when compromised lead to major pathologies.

