## Supplemental figures for "Direct interactions of CEACAM1 and CD36 with micellar LPS and each other"

**Supplemental Figure S1. Expression of sCEACAM1-4D**. **A**. Amino acid sequence of construct showing leader sequence (grey), N-domain (red), A1 domain (green), B1 domain (blue), A2 domain (purple) and interdomain sequences (black). **B.** SEC analysis of purified protein (bottom), SEC analysis of anti-N-domain antibody T84.1 (middle) and co-incubation of sCEACAM1-D with T84.1 antibody (top).

**A**

MGHLSAPLHRVRVPWQGLLLTASLLTFWNPPTTAQLTTESMPFNVAEGKEVLLLVHNLPQQLFG
YSWYKGERVDGNRQIVGYAIGTQQATPGPANSGRETIYPNASLLIQNVTQNDTGFYTLQVIKSD
LVNEEATGQFHVYPELPKPSISSNNSNPVEDKDAVAFTCEPETQDTTYLWWINNQSLPVSPRLQ
LSNGNRTLTLLSVTRNDTGPYECEIQNPVSANRSDPVTLNVTYGPDTPTISPSDTYYRPGANLS
LSCYAASNPPAQYSWLINGTFQQSTQELFIPNITVNNSGSYTCHANNSVTGCNRTTVKTIIVTE
LSPVVAKPQIKASKTTVTGDKDSVNLTCSTNDTGISIRWFFKNQSLPSSERMKLSQGNTTLSIN
PVKREDAGTYWCEVFNPISKNQSDPIMLNVNYNALPQEN

**B**

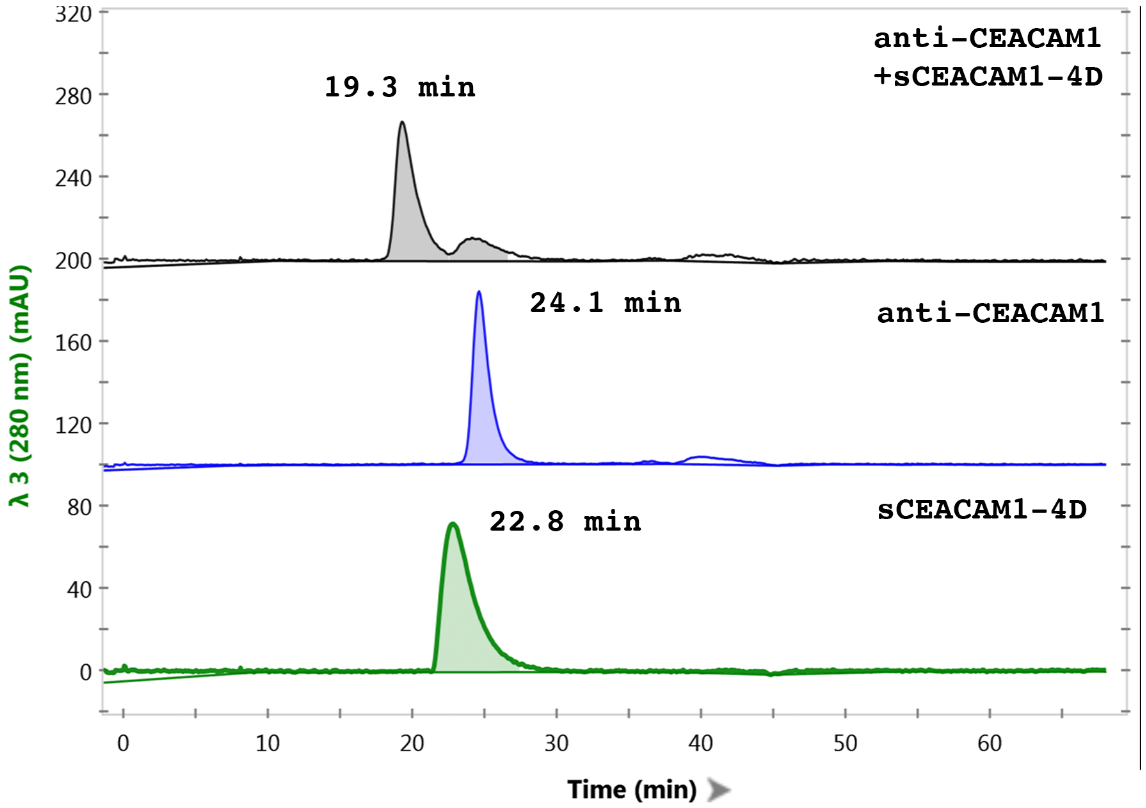

**Supplemental Fig S2. Steady state affinity binding of LPS-Ra to sCEACAM1-4D, CEACAM1-N domain and sCD36. A.** sCEACAM-4D. **B.** CEACAM1-N domain. **C**. sCD36. All data points in duplicate. Chi square values for **A-C** are 2.55, 2.63, and 0.56, respectively.

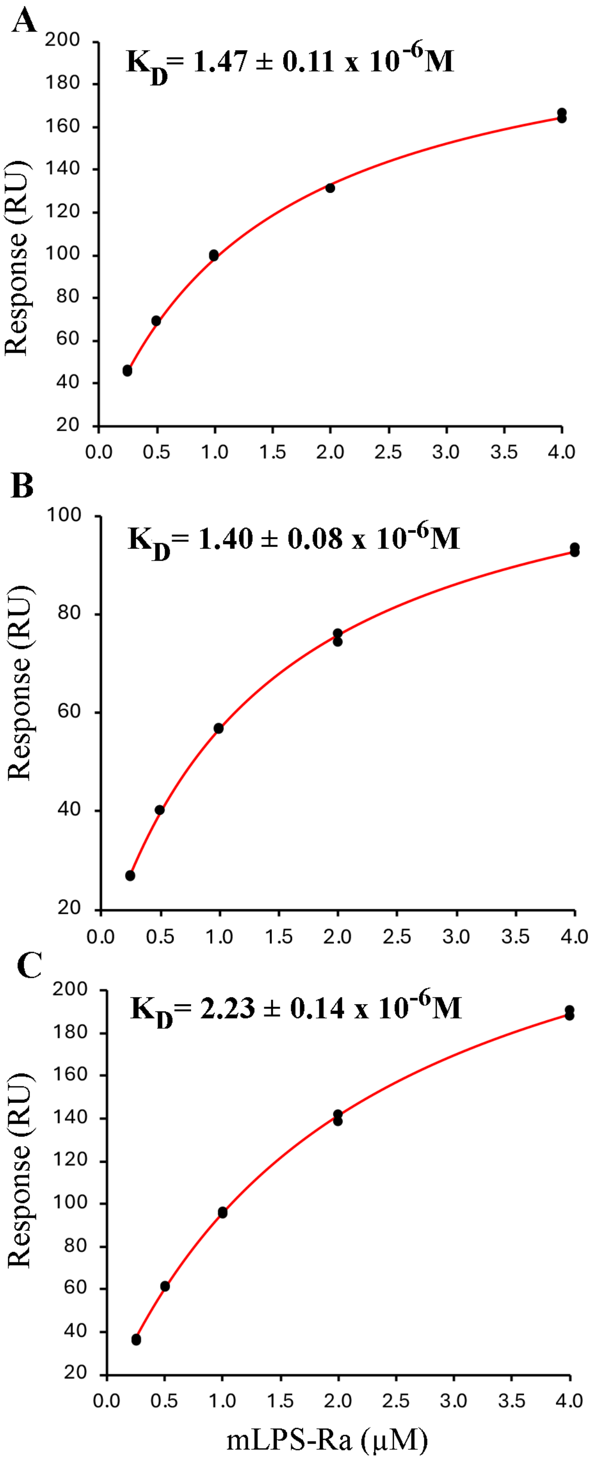

**Supplemental Fig S3. LPS-Ra binding to sCEACAM1-4D vs CEACAM1-N domain. A.** Continuous injection (180 s) of LPS (0.5 µM) to immobilized sCEACAM1-4D was followed by regeneration with 0.5% SDS. Then continuous injection (180 s) of anti-CEACAM1 antibody T84.1 (0.15 µM) was followed by continuous injection (180 s) of LPS (0.5 µM). RUs bound at each step are shown. **B**. Purification of expressed CEACAM1-N (N-domain with His6 tail) showing amino acid sequence (signal sequence in black, coding sequence in green, His6 in red) followed by SEC and SDS gel analysis of purified product. **C**. Binding of LPS-$R_a$ to immobilized CEACAM1-N.

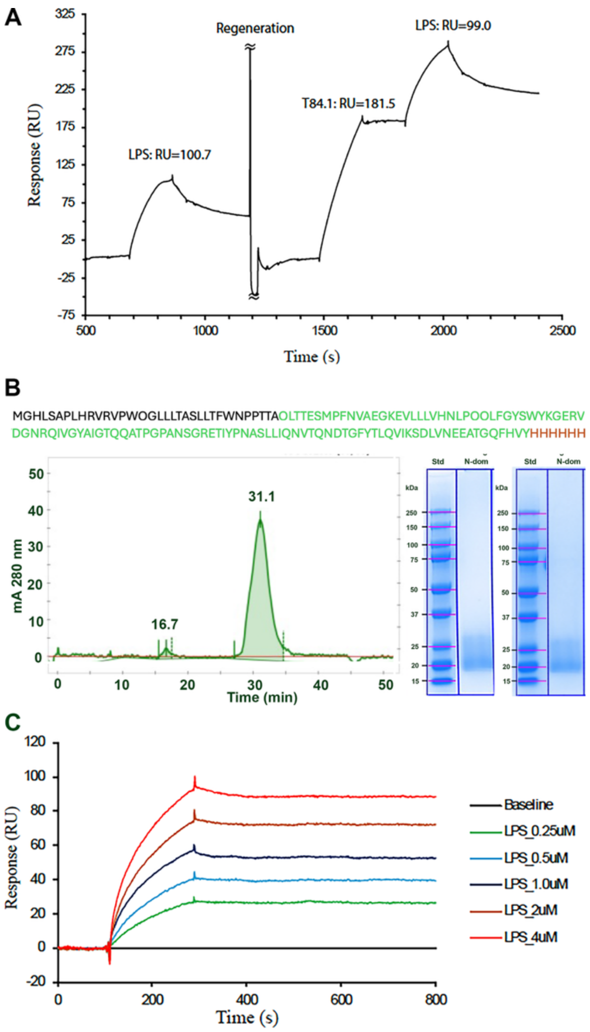

**Supplemental Fig S4. Plot of normalized binding (RU) of LPS with increasing ratios of bile acids cholate or deoxycholate to immobilized sCD36 or sCEACAM1-4D.** The LPS concentration is 2 μM, and the concentration of bile acids (cholate and deoxycholate) are 24, 48, 96, 192 and 384 μM. The RUs from LPS (2 μM) in absence of bile acids is set to 1.0. The RU from LPS and bile acids mixtures is normalized against RUs of free LPS (2 μM). **A.** Normalized SPR RUs of LPS binding to CD36 versus the molar ratio of bile acids/LPS. LPS_Ch and LPS_DC are RUs from LPS in the presence of cholate or deoxycholate; the LPS_Ch-Ch and LPS_DC-DC are the RU of LPS and BA mixture subtracting RU of free cholate or deoxycholate. **B**. Normalized SPR RUs of LPS binding to Ceacam1 versus the molar ratio of bile acids/LPS. The definitions of curves are the same to (**A**) using CEACAM1-4D as the immobilized ligand.

**A**

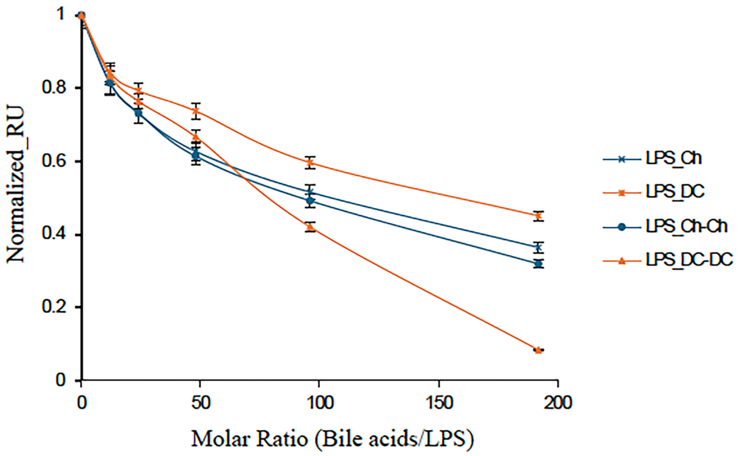


**B**

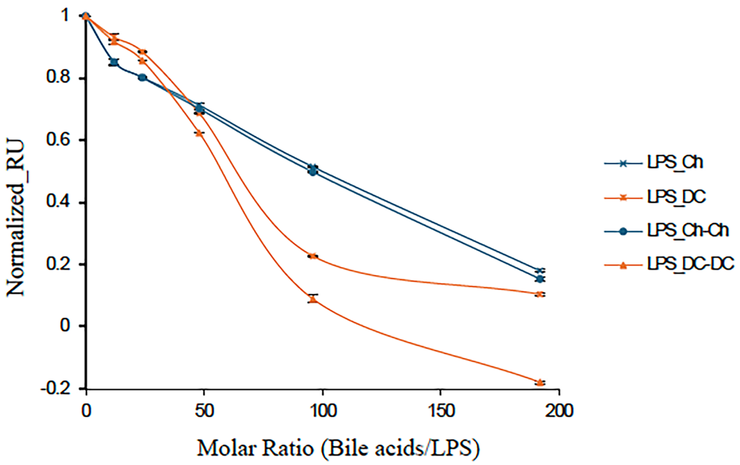

**Supplemental Figure S5. Calculation of Stokes radii for sCD36 and sCEACAM1-4D. A.** Plot of standards (log10 Rs) vs Kd. **B.** Calculations for standards. **C.** Calculations for CD36 and CEACAM1-4D and for each with LPS-Ra.

A

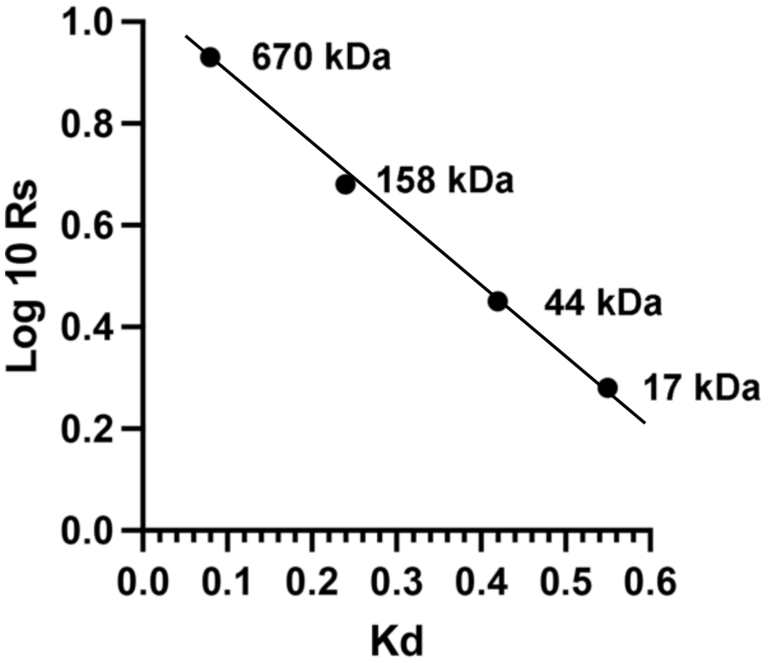


B

| MW | Ve | Ve-Vo | (Ve-Vo)/(Vt-Vo)= Kd | Lit Rs (nm) | Log10 Rs |
|---|---|---|---|---|---|
| 670 kDa | 9.0 mL | 1.5 mL | 0.08 | 8.6 | 0.93 |
| 158 kDa | 11.8 mL | 4.3 mL | 0.24 | 4.8 | 0.68 |
| 44k Da | 14.9 mL | 7.4 mL | 0.42 | 2.8 | 0.45 |
| 17kDa | 17.2 mL | 9.7 mL | 0.55 | 1.9 | 0.28 |

C

| peak | Ve | Ve-Vo | (Ve-Vo)/(Vt-Vo)= Kd | Log 10Rs | Rs (nm) |
|---|---|---|---|---|---|
| CD36 | 13.1 mL | 5.6 mL | 0.32 | 0.60 | 4.0 |
| CD36 +LPS | 9.5 mL | 2.0 mL | 0.11 | 0.90 | 7.9 |
| CEACAM1-4D | 11.2 mL | 3.7 mL | 0.21 | 0.74 | 5.5 |
| CEACAM1-4D +LPS | 8.3 mL | 0.8 mL | 0.045 | 0.95 | 8.9 |

**Supplemental Fig S6. Amino sequences of CD36-BioID2 fusion protein and CEACAM1-4L.** Sequence of CD36-BioID2 fusion protein that was inserted in plasmid 25ABWPVD-4062235 from Addgene. Sequence of human CD36 in green and BioID2 from *Aquifex aeolicus* (biotin-[acetyl-CoA-carboxylase] ligase) in red, cloning introduced sequences in blue.

MGCDRNCGLIAGAVIGAVLAVFGGILMPVGDLLIQKTIKKQVVLEEGTIAFKNWVKTGTEVYRQ
FWIFDVQNPQEVMMNSSNIQVKQRGPYTYRVRFLAKENVTQDAEDNTVSFLQPNGAIFEPSLSV
GTEADNFTVLNLAVAAASHIYQNQFVQMILNSLINKSKSSMFQVRTLRELLWGYRDPFLSLVPY
PVTTTVGLFYPYNNTADGVYKVFNGKDNISKVAIIDTYKGKRNLSYWESHCDMINGTDAASFPP
FVEKSQVLQFFSSDICRSIYAVFESDVNLKGIPVYRFVLPSKAFASPVENPDNYCFCTEKIISK
NCTSYGVLDISKCKEGRPVYISLPHFLYASPDVSEPIDGLNPNEEEHRTYLDIEPITGFTLQFA
KRLQVNLLVKPSEKIQVLKNLKRNYIVPILWLNETGTIGDEKANMFRSQVTGKINLLGLIEMIL
LSVGVVMFVAFMISYCACRSKTIKGARRFKNLIWLKEVDSTQERLKEWNVSYGTALVADRQTKG
RGGLGRKWLSQEGGLYFSFLLNPKEFENLLQLPLVLGLSVSEALEEITEIPFSLKWPNDVYFQE
KKVSGVLCELSKDKLIVGIGINVNQREIPEEIKDRATTLYEITGKDWDRKEVLLKVLKRISENL
KKFKEKSFKEFKGKIESKMLYLGEEVKLLGEGKITGKLVGLSEKGGALILTEEGIKEILSGEFS
LRRSYPYDVPDYA
